# Landscape of transcript isoforms in single T cells infiltrating in non-small cell lung cancer

**DOI:** 10.1101/2020.01.29.924308

**Authors:** Jiesheng Li, Zemin Zhang, Xianwen Ren

## Abstract

Single cell RNA-seq has enabled high-resolution characterization of molecular signatures of tumor-infiltrating lymphocytes. However, analyses at the transcript isoform level are rarely reported. As alternative splicing is critical to T cell differentiation and activation, here we proposed a computational method named as IDEA to comprehensively detect and annotate differentially used isoforms across cell subtypes. We applied IDEA on a scRNA-seq dataset of 12,346 T cells from non-small cell lung cancer. We found most genes tend to dominantly express one isoform in single T cells, enabling typing T cells according to the isotypes given a gene. Isotype analysis suggested that tumor-infiltrating T cells significantly preferred specific isotypes for 245 genes in CD8+ T cells and 456 genes in CD4+ T cells. Functional annotation suggests that the preferred isoforms involved in coding/non-coding switches, transcription start site changes, gains/losses of domains and subcellular translocation. Clonal analysis revealed that isoform switching occurred during T cell activation/differentiation. Our analysis provides precise characterization of the molecular events in tumor-infiltrating T cells and sheds new lights into the regulatory mechanisms of tumor-infiltrating T cells.

## INTRODUCTION

Tumor-infiltrating T cells play important roles in antitumor immunity. Immunotherapies targeting T cells, e.g., immune checkpoint blockade, have shown promising therapeutic effects in multiple cancer types (1). However, the outcomes of these therapies are inconsistent across patients, of which the complex microenvironment within tumors may have critical impacts. Analysis of tumor-infiltrating T cells by single cell RNA sequencing (scRNA-seq) may provide critical clues for the mechanisms underlying resistance to immunotherapies by facilitating a detailed survey of T cell subtypes and molecular characteristics in the tumor microenvironment. Tumor-infiltrating T cells have been characterized by scRNA-seq in multiple cancer types including liver (2), non-small-cell lung(3), and colorectal cancer(4), breast carcinoma(5), head and neck carcinoma(6), and melanoma(7). However, almost all the current studies focused only on the gene-level characterization, with the transcript-level analysis rarely reported.

T cells utilize differential isoforms to regulate their functions throughout lifetime. The ImmGen Consortium integrated high-throughput data and revealed pervasive events of alternative splicing (AS) in T cells, with ~60% of genes showing different AS events linking to the differentiation of different T cell lineages(8). Multiple studies identified specific transcript isoforms involved in T cell-related biology(9), including T cell activation (10–12), T cell differentiation(13, 14), and apoptosis(15). For instance, *PTPRC*, which encodes the transmembrane tyrosine phosphatase CD45, is critical for T cell receptor (TCR) signal transduction(16). Naive T cells express long isoforms for *PTPRC*, with one or more of the alternative exons (No. 4,5, and 6) retained(16, 17). While in memory T cells, short isoforms with the alternative exons (No. 4,5, and 6) skipped are expressed for *PTPRC*(16, 17). Another example gene is PDCD1 encoding the programmed cell death protein 1 (PD-1), which down-regulates the T cell activity and promotes self-tolerance via receiving the signals of PD-L1 or PD-L2(18). A short isoform with 42-nucleotide in-frame deletion of *PDCD1* is found expressed in peripheral blood mononuclear cells (PBMCs), of which the encoded protein does not engage PD-L1/PD-L2 and can induce proinflammatory cytokines via a recombinant form(19). These examples together with the pervasive alternative isoforms identified in T cells suggested the essential roles of alternative splicing events in the development and functional properties of T cells and the necessity of isoform-level characterization. However, it remains unknown that how T cell subtypes utilize isoforms at single-cell resolution in tumors.

Here, we established a computational pipeline, named as IDEA, to investigate the single cell transcript landscape based on scRNA-seq data. By analyzing the tumor-infiltrating T cells in non-small cell lung cancer patients(3), we found that genes with multiple isoforms tend to dominantly express one isoform in individual T cells, allowing the definition of isotypes for each cells to characterize the preferred isoform usage. Enrichment analysis based on the isotypes of individual T cells suggested that for hundreds of genes T cell subtypes exhibit significantly differential preferences for isotypes, particularly for the tumor-infiltrating exhausted CD8+ and regulatory CD4+ T cells. The different preferences of isotypes between T cells subtypes lead to distinct biological functions including coding/non-coding translocation. Clonal analysis based on TCR sequences reveals that the differential preference occurred via isoform switching, which is a pervasive process during T cell clonal expansion. Altogether, our work provides a comprehensive analysis of the transcript isoform landscape of tumor-infiltrating T cells in non-small cell lung cancer at the single-cell resolution. The abundant isoform switching events identified among tumor-infiltrating T cell subtypes may serve as a rich data resource for precisely analyzing the mechanisms underlying T cell-based antitumor immunity. Our analysis also highlights the necessity of splicing analysis at the single-cell resolution, which can be applied to a wide range of biological questions.

## MATERIAL AND METHODS

### ScRNA-seq dataset used in this study

We used the scRNA-seq dataset generated by Guo et al.(3) to investigate differential isoform usages of tumor-infiltrating T cells at the single cell level. Corresponding bulks were also downloaded. This dataset contains 12,346 T cells from 14 treatment-naive non-small-cell lung cancer patients. All the T cells were isolated through FACS using canonical T cell markers CD3D, CD8A, CD4, CD25, and the viability staining solution 7AAD. On these T cells, single-cell transcriptome amplification were performed following Smart-seq2 protocol(20). The resulting cDNA libraries were sequenced by Illumina Hiseq 2500 sequencer (100 bps, pair-end) or Illumina Hiseq 4000 sequencer (150 bps, pair-end) at an average depth of 1.04 million uniquely-mapped read pairs per cell without obvious 3’ bias along gene bodies. Such quality of data thus enabled the detection of expressed isoforms reliably. tSNE plots are obtained as described in the original publication using gene level expression. The cell type labels for each T cell were also obtained and the corresponding properties of them were presented in the table below.

| T cell cluster name | Presumptive functional properties |
| --- | --- |
| CD8-C1-LEF1 | Naive CD8+ T cells |
| CD8-C2-CD28 | - |
| CD8-C3-CX3CR1 | Effector CD8+ T cells |
| CD8-C4-GZMK | - |
| CD8-C5-ZNF683 | Tissue resident memory CD8+ T cells |
| CD8-C6-LAYN | Exhausted CD8+ T cells |
| CD4-C1-CCR7 | Naive CD8+ T cells |
| CD4-C2-ANXA1 | Central memory CD4+ T cells in blood |
| CD4-C3-GNLY | Effector CD4+ T cells |
| CD4-C4-CD69 | - |
| CD4-C5-EOMES | - |
| CD4-C6-GZMA | - |
| CD4-C7-CXCL13 | Exhausted CD4+ T cells |
| CD4-C8-FOXP3 | Resting CD4+ T cells |
| CD4-C9-CTLA4 | Suppressive CD4+ T cells |

### Overview of IDEA for isoform analysis at single-cell resolution

ScRNA-seq has been widely applied in biomedical studies and resulted in revolutionary discoveries in various fields including cell development and cancer immunology (2–7). However, the current analysis is mainly gene-focused, with the transcript isoforms seldom depicted. To identify isoform expression at single-cell resolution, characterize the preference by distinct cell subtypes, and illustrate the potential biological impacts, we proposed a computational method, named as IDEA (Isoform Detection, Enrichment and functional Annotation), based on full-length scRNA-seq data (**Figure 1**). Basically, the framework of IDEA is composed by three components: 1) isoform quantification to detect the dominant isoforms expressed by individual cells; 2) enrichment analysis of cells dominantly expressing specific isoforms across cell types; and 3) functional annotation of isoforms differentially expressed by cells from different cell types. For cells that can trace the developmental lineages, e.g., T cells by T cell receptors, IDEA also provides a functional module to detect genes undergoing isoform switching. The technical details of IDEA were stated as follows.

**Figure 1.**
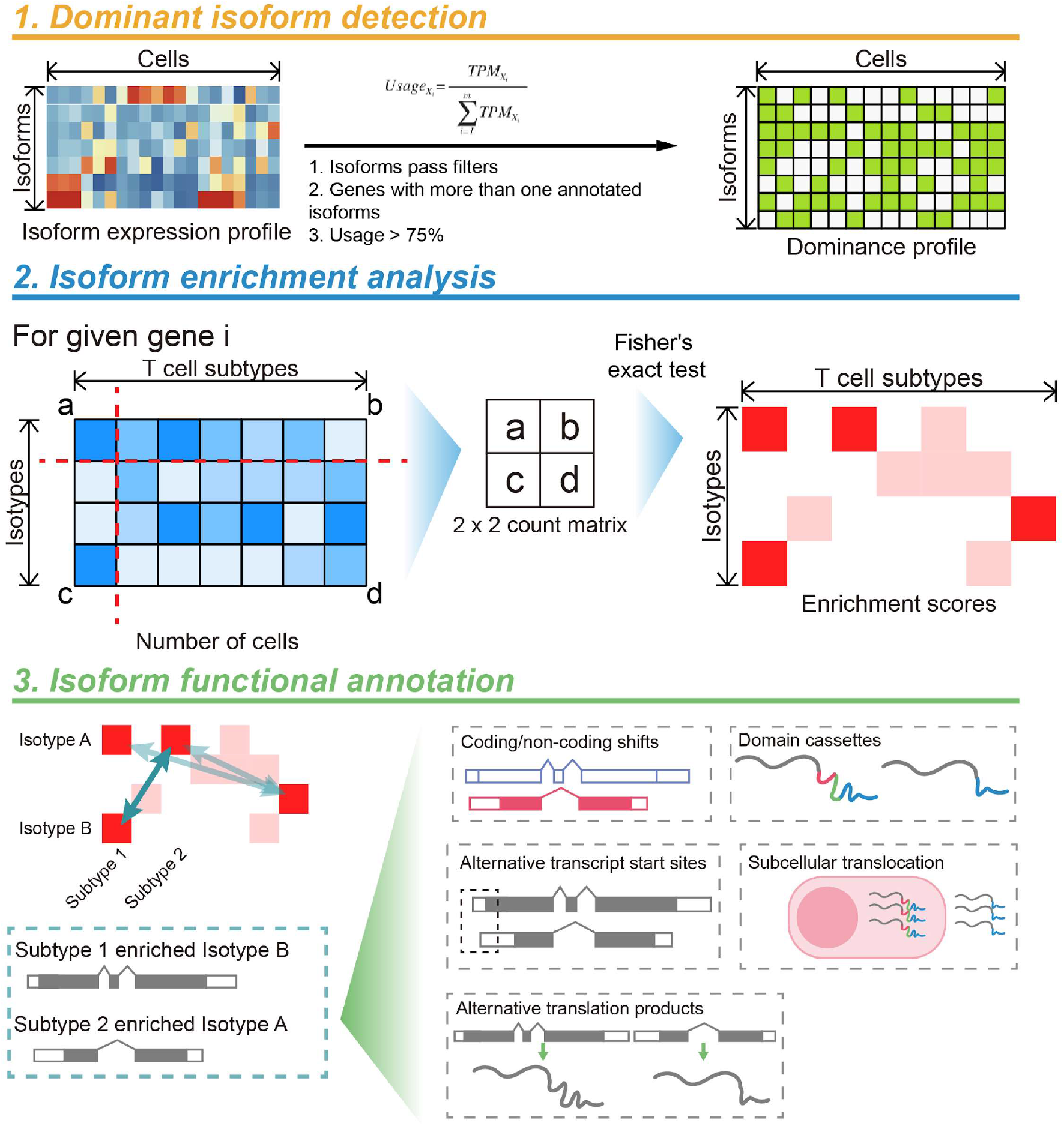
Overview of IDEA for single-cell isoform analysis.

### Quantification of the expression level and isoform usage of transcripts

First, STAR(21) with the default parameters was used to align scRNA-Seq data to the human genome sequences (hg38, available from https://www.gencodegenes.org/human/). Stringtie(22) with parameter “-eB” was then used to parse the aligned “bam” files with the input of the gene annotation gtf file (the same version with the genome sequences, from GENCODE). The python script “prepDE.py” from the Stringtie website with parameter “-l 100” was used to estimate the expression levels of transcript isoforms. Batch effects among samples were removed by Pagoda2 (https://github.com/hms-dbmi/pagoda2). The TPM (transcripts per million) table for isoforms was then derived from the adjusted count table by normalizing the effects of transcript length using self-implemented function. The expression levels for genes were calculated as the sum of the corresponding isoforms. Transcripts expressed (adjusted count > 0) in less than 30 cells were filtered from downstream analysis. Cells with the number of expressed transcripts (adjusted count > 0) less than 1,000 were filtered from the downstream analysis. Outlier analysis using the ‘isOutlier’ function implemented by the R package scater (23) was applied to the following features: 1) number of expressed transcripts with parameters ‘nmads=3, type=“lower”, log=TRUE’; 2) counts of mitochondrial genes with parameters ‘nmads = 3, type = “lower”, log = F’; and 3) number of uniquely mapped reads with parameters ‘nmads = 3, type = “lower”, log = T’. Cells that were identified as outliers or without cluster annotated in the original paper (indicating low-quality or doublets) were filtered out․.11,595 out of 13,154 cells were retained for the downstream analysis.

After filtering, the transcripts per million (TPM) table for isoforms was derived from the adjusted count table considering the different length of each transcript. Gene expressions were calculated as the sum of its corresponding isoforms. When the expression levels of transcript isoforms were determined, given a gene, the isoform usages were calculated by the following formula:

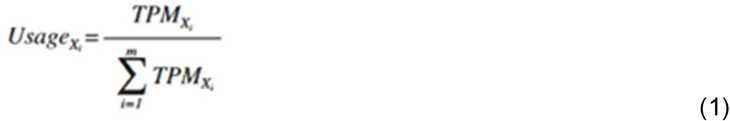

where was the TPM value of the *i^th^* isoform of gene *X* in a given sample, and m is the total number of transcript isoforms of gene *X* in the GENCODE annotation. The isoform usage is naturally between 0 and 1.

### Evaluating the dominance of isoforms in single cells

We devised two metrics to quantitatively evaluate the dominance of isoform expression across genes in single cells, i.e, entropy and H-index. For each cell, we discretized the isoform usages of all transcripts into 100 bins. The fraction of isoforms in the i^th^ bin is denoted as *p*_*i*_. We used the following formula to estimate the entropy of isoform usage for the given cell:

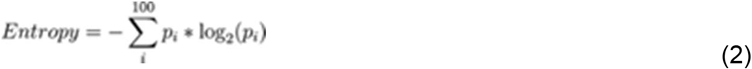

Entropy measures the evenness of the isoform usage distribution, with a high entropy value indicating a close-to-uniform distribution between 0 and 1. Dominant expression of one isoform for most genes would result in low entropy values. The second metric was H-index. H-index is initially a bibliometrics measure to evaluate both the productivity and citation impact of a scholar. Here we borrowed this measure to quantify the frequency of dominant isoforms in a cell, which is defined by the following formula:

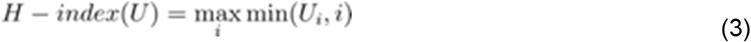

Where *i* is the normalized index of a transcript in all expressed transcripts of all genes sorted in a descending manner, *u*_*i*_ is the corresponding usage in the given cell, and the H-index gives out the fraction of isoforms that have the corresponding usage more than the same value. For example, a cell with H-index 0.7 means at least 70% of expressed isoforms in this cell have usage that exceed 0.7. Hence higher the H-indexes indicates more genes having dominant isoforms expressed.

We constructed pseudo-bulk RNA-seq data by merging all TTCs from patients, P1202 and P1010 individually, and collected the real bulk RNA-seq data of P1010. A series of sequencing depth were simulated by random sampling reads from merged bulks and real bulks using seqtk (https://github.com/lh3/seqtk), in orderto evaluate the impacts of sequencing depth on the dominance of isoforms. The same isoform quantification pipeline was applied to all these bulk RNA-seq data.

### Isotypes of single cells

The dominant expression of one isoform in single cells allows defining the isotypes of single cells when the gene is given. To analyze the isotypes of single T cells in lung cancer and reach robust conclusions, we excluded those low expressed genes that had TPMs of all the isoforms < 3. Given a cell, the isoform of one gene with the usage > 0.75 was defined as the dominant isoform in this cell, and the isotype of this cell for the given gene was labeled as “T_GENE_^ISOFORM^”. For genes that had no isoform with usage > 0.75, the isotype was labeled as T_GENE_^MIX^. Here we used 0.75 as the threshold to identify the dominant isoform, which can guarantee that the usage of the top isoform is at least 3-fold of the second isoform. More stringent cutoffs can also be used but will not change the observations in this study. A virtual isotype labelled as T_GENE_^NO_EXPR^ was introduced to denote the phenotypes of single cells that did not express the specific genes.

### Quantifying the enrichment of different isotypes in T cell subtypes

Given a gene, we applied Fisher’s exact test to examine whether an isotype was enriched in a specific T cell subtype. CD8+ and CD4+ T cells were analyzed separately because of their distinct developmental trajectories. Given an isotype (target isotype), we categorized all the CD8+ or CD4+ T cells into 4 categories by examining whether a cell is labeled as the target isotype and whether the cell belongs to the target T cell subtype. Then, a 2 × 2 contingency table was constructed and used for Fisher exact test. Since each gene with multiple isoforms would be tested multiple times for every possible isotype-subtype combination, the resulting p values were then adjusted based on the Benjamini-Hochberg method for multiple testing correction, and the enrichment scores are defined as −log_10_ (adjusted p values). Each enrichment score that exceeds 1.3 (adjusted p-value < 0.05) was used to identify significantly enriched isotypes in a given cell cluster.

### Functional annotations of the enriched isoforms

Functional annotations of the transcript isoforms including transcript types and transcription start sites were extracted from ENSEMBL(24). Transcript types were further re-categorized according to the biotypes for comprehensive analysis according to ENSEMBL (https://asia.ensembl.org/Help/Faq?id=468). For coding isoforms, we used InterPro(25) to acquire protein structural information including signal peptides, transmembrane region and Pfam domains. Furthermore, we combined signal peptides and transmembrane regions to determine the potential subcellular locations. Proteins with at least one transmembrane region were assumed to be membrane-integrated. Proteins without any transmembrane region were examined for the existence of the signal peptides. If the signal peptides exist, the protein were assumed to be secreted. Otherwise, the protein were assumed to be cytoplasmic.

### Identification and categorization of differential isotypes

For a given gene, differential isotype usages were defined as a pair of isotypes enriched in two cell subtypes. The same isotypes that were enriched by both clusters are excluded. The differential isotype usages were then examined for the coding potentials, transcription start sites and protein domains of the isoforms involved and thus categorized into 5 categories: coding/non-coding switches, transcription start site changes, gains/losses of domains and subcellular location changes. That is, for each pair of differential isotypes, if they have different coding potentials (e.g. one of them is annotated as ‘coding’, the other one is annotated as ‘non-coding’), then this pair fall in the category of coding/non-coding switches. The similar comparison was conducted for all the features mentioned above.

### Pathway analysis

Both Gene Ontology (GO) term enrichment analysis and pre-ranked gene set enrichment analysis (GSEA) were performed using an R implementation (package clusterProfiler)(26) on genes with differential isotypes. Specifically, genes for GSEA were sorted by enrichment scores in a decreasing order.

### Clonal analysis

Clonotypes of T cells were acquired as previously described(3). That is, the TCR sequences were assembled and quantified by TraCeR(27) and dominant alpha-beta pairs were identified for each T cell. Each unique, productive (i.e., in frame) and dominant alpha-beta pair was defined as a clonotype and cells harboring the same clonotype would be considered clonal. Among the cells passed filters described above, we identified 659 clonotypes were shared by at least two T cells.

Within each clonotype, we examined the number of isotypes for each gene. Genes with at least two isotypes expressed in at least one clonotype would be assumed to undergo isoform switching since the expression pattern was likely to switch between isotypes during the clonal expansion. P values for the statistical significance of the overlap between genes with isoform switching and genes with differential isotypes across T cell subtypes was evaluated by assuming a hypergeometric distribution using R function phyper with parameter ‘lower.tail = FALSE’.

## RESULTS

### The presence of dominant isoforms in single cells compared with bulk tissues

To comprehensively investigate differential isoform preferences among T cell clusters, we first perform isoform quantification to obtain the basic properties of single cells at isoform level. Saturation analysis indicated that with proper cutoff for expression, we could reliably detect functional classes of isoforms, including cytokines and transcription factors (**Supplementary Figure. 1a**). The usages of all expressed isoforms from both T cells and bulks are calculated (**See methods**) and displayed distinct bimodal distribution (**Figure 2a**). Comparison of the usage distribution of all expressed isoforms between single cell samples and the corresponding bulk sample shows that single cells utilize isoforms more dominantly (**Figure 2b**), which means that single cells tend to dominantly express one isoform while others remain low or have no expression. This phenomenon is pervasive when comparing other single cells and bulk samples (**Figure 2c**) via two quantitative indexes, entropy and H-index (**See methods**). Entropy, an index indicating the “unevenness” of the isoform expression distribution with lower values, showed that isoform expression in single cells was significantly skewed when compared to bulks, while H-index, an author-level metric often used to quantify the productivity and citation impact of a scholar, showed that single cells had more genes that were dominantly expressing one isoform. However, due to different sequencing depths between single cells and bulk samples, we could not affirm that the exhibited dominant isoform expressions are inherent properties of single cells based on this analysis. Therefore, we constructed a series of compositional bulks with down-sampled reads from real bulk samples provided by Guo and bulks of merged T cells (**See methods**). We found that, even at the same sequencing depth, single cells still had significantly lower entropies (< 3) compared to subsampled bulks (4–5), and higher H-indexes (around 0.8) compared to H-indexes of subsamples ranging from 0.5 to 0.7 (**Figure 2d,e**). H-indexes of compositional bulks decreased while sequencing depth increased. Inversely, entropy increased with sequencing depth, suggesting that fewer genes are exhibiting dominant expression of one isoform at higher sequencing depths in compositional bulk samples. Thus we confirmed that single T cells tend to be dominated by one of many isoforms of a multi-isoform gene, which has also been reported in previous studies based on mouse brain and simulation data(28, 29).

**Figure 2.**
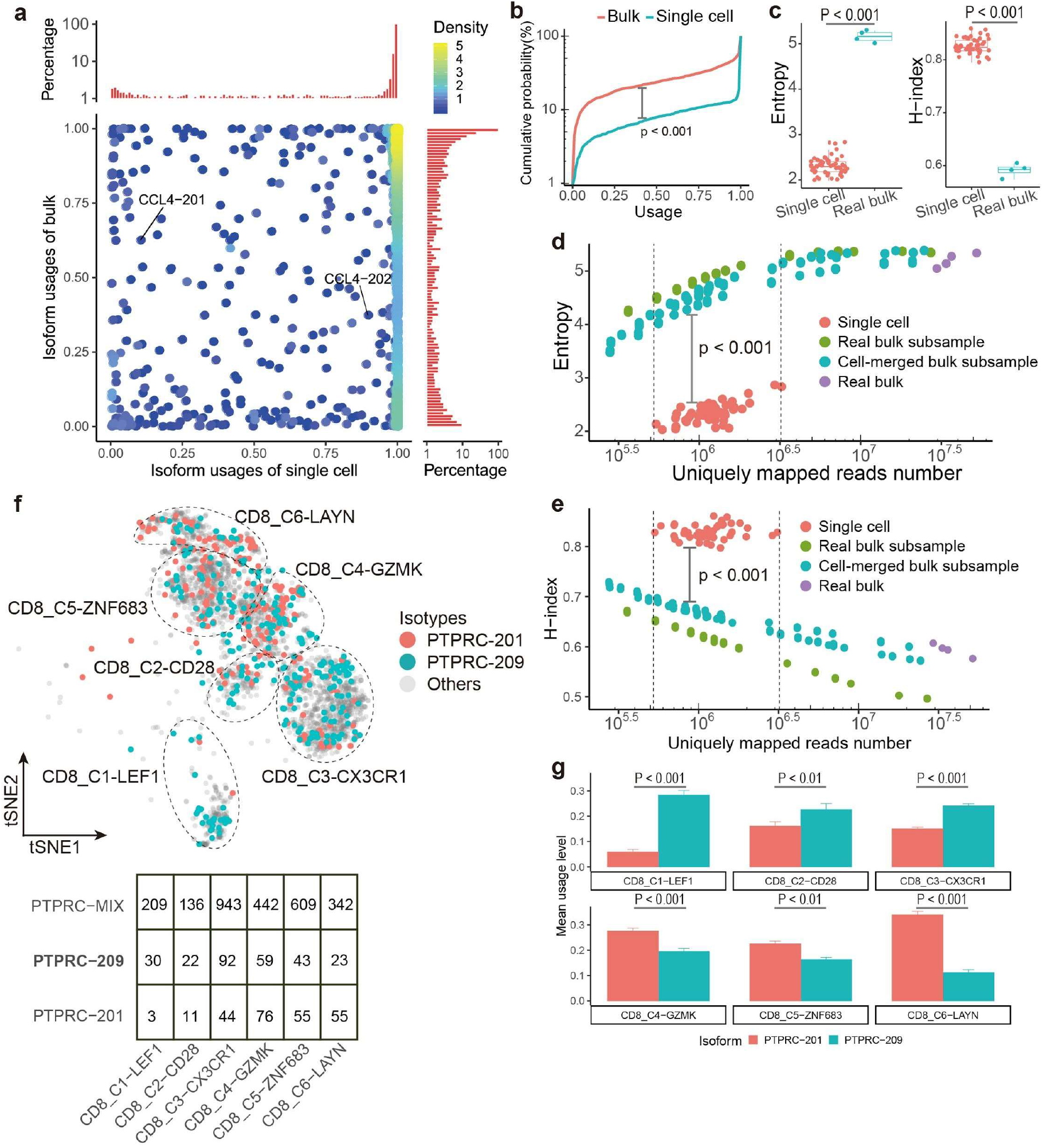
Single cells have a distinct isoform usage distribution from bulk tissues. **a**, comparison of the isoform usage distributions of the bulk sample P1010T and the single cell TTC34-1010. Each dot denotes a transcript isoform and the usage differences between single cells and bulk tissues are exemplified by CCL4. Y-axes (top and right bar plots) represent the relative frequency of each usage bin (normalized with the highest bar). Bar plots display the distribution for each sample correspondingly, with y-axis represent the percentage of each given bin of usages compared with the highest bar. **b**, the cumulative distributions for samples in **a** show statistically significant differences (KS test). **c**, quantitative assessment by entropy (left) and H-index (right) reveals that the distinct isoform usage distributions observed in **a** and **b** can be extended to other single cells and their paired bulk tissues. Each dot denotes a single cell or bulk sample. Low entropy (**d**) and high H-index (**e**) of single cells compared with bulk samples are not caused by different sequencing depth, where “pseudo-bulk by subsampling” means pseudo-samples constructed by subsampling reads of real bulk samples and “pseudo-bulk by merging” means pseudo-samples constructed by merging different single cells. **f**, dominant isoform usage in single cells allows the identification of “isotypes” of single cells for each gene with multiple transcript isoforms (exemplified by the CD8+ T cells and the gene *PTPRC*). Cell counts of each isotypes were recorded in the table shown below. **g**, isotypes of *PTPRC* show different enrichment in T cell clusters defined by gene expression profiles, where PTPRC-209-dominating cells are significantly enriched in blood-origin T cell clusters (top) while PTPRC-201-dominating cells are enriched in tissue/tumor-origin T cell clusters (bottom). Wilcoxon test is used to evaluate the statistical significance, error is s.e.m.

To specify the dominant isoform for each gene, we labeled cells as “isotypes” (**See methods**). As a proof of the concept, we investigated *PTPRC*, one of the best-studied examples, to examine the isotypes in T cells. *PTPRC* is essential during lymphocyte differentiation, using exons 4, 5, and 6 variably(13, 30). Two transcript isoforms are highlighted according to ENSEMBL(24): PTPRC-209, which retains exons 4,5,6 and translates into protein isoform PTPRC-RABC, and PTPRC-201, which splices out all three exons and translates into PTPRC-RO (**Supplementary Figure 1b**). PTPRC-RABC and PTPRC-RO are known to be expressed by distinct T cell populations; the expression of RA (isoforms with exon 4) is associated with the naive state while memory cells switch to RO completely at the protein level(30, 31). Thus we focus mainly on PTPRC-209 and PTPRC-201 in the following analysis.

Although no obvious differences were observed on the gene level across T cell subtypes in our dataset (**Supplementary Figure 1c**), we identified various isotypes of PTPRC in each T cell subtype, including T_PTPRC_^201^, T_PTPRC_^209^ and T_PTPRC_^MIX^ (**Figure 2f**). As expected, more PTPRC-209-dominated cells were counted among naive CD8+ T cells (CD8_C1-LEF1) and more PTPRC-201-dominated cells were among exhausted CD8+ T cells (CD8_C6-LAYN). Furthermore, the mean usage level of isoform PTPRC-209 is significantly higher than PTPRC-201 in naive T cells (CD8_C1-LEF1), CD8_C2-CD28, and effector T cells (CD8_C1-LEF1), while the mean usage level of PTPRC-201 is significantly higher in “pre-exhausted” CD8+ T cells (CD8_C4-GZMK), tissue resident memory CD8+ T cells (CD8_C5-ZNF683), and exhausted T cells (CD8_C6-LAYN) (**Figure 2g**). The inverse usage relationship of these two isoforms in CD8+ T cell subtypes coincides with the current knowledge(30, 31). Thus, our results not only confirm the preferences for *PTPRC* isoforms across T cell subtypes, but also illustrate that PTPRC-201 and PTPRC-209 were expressing in an inverse way.

### Different T cell subtypes prefer to express different isotypes for the same genes

To further investigate the preferences of T cell subtypes for certain isotypes, we performed enrichment analysis for all of the multi-isoform genes (**See methods**). Since CD4 and CD8+ T cells differ in cell types, we performed enrichment analysis on them separately. For CD8+ T cells, 5323 out of 14629 genes have at least one isotype enriched in any T cell subtypes, and 557 of them have multiple isotypes enriched in at least two CD8+ T cell subtypes (**Supplementary Figure 2a**). As for CD4+ T cells, the number of genes with enriched isotypes was slightly more than that of CD8+ T cells (6241 genes), and about 17% (1044/6241) of genes with multiple enriched isotypes were detected in more than one CD4+ T cell type (**Supplementary Figure 2b**).

CD8 exhausted T cells were found to derive the most genes (3650 genes) with significant enriched isotypes (**Figure 3a**). This phenomenon is not influenced by the number of cells in each cell subtype (**Supplementary Figure 2c**), which might suggest that CD8 exhausted T cells are in distinct states on isoform level compared with other CD8+ T cell subtypes. Furthermore, Gene ontology (GO) enrichment analysis reveals that the genes detected in exhausted T cells are involved in mRNA processing, cell cycle, antigen processing and presentation, T cell receptor signaling, and response to hypoxia (**Figure 3b**), while gene set enrichment analysis (GSEA) suggests that isoforms are utilized in various pathways of exhausted T cells, such as T cell activation, T cell receptor signaling pathway and response to hypoxia (**Figure 3c**; **Supplementary Figure 2e,f**).

**Figure 3.**
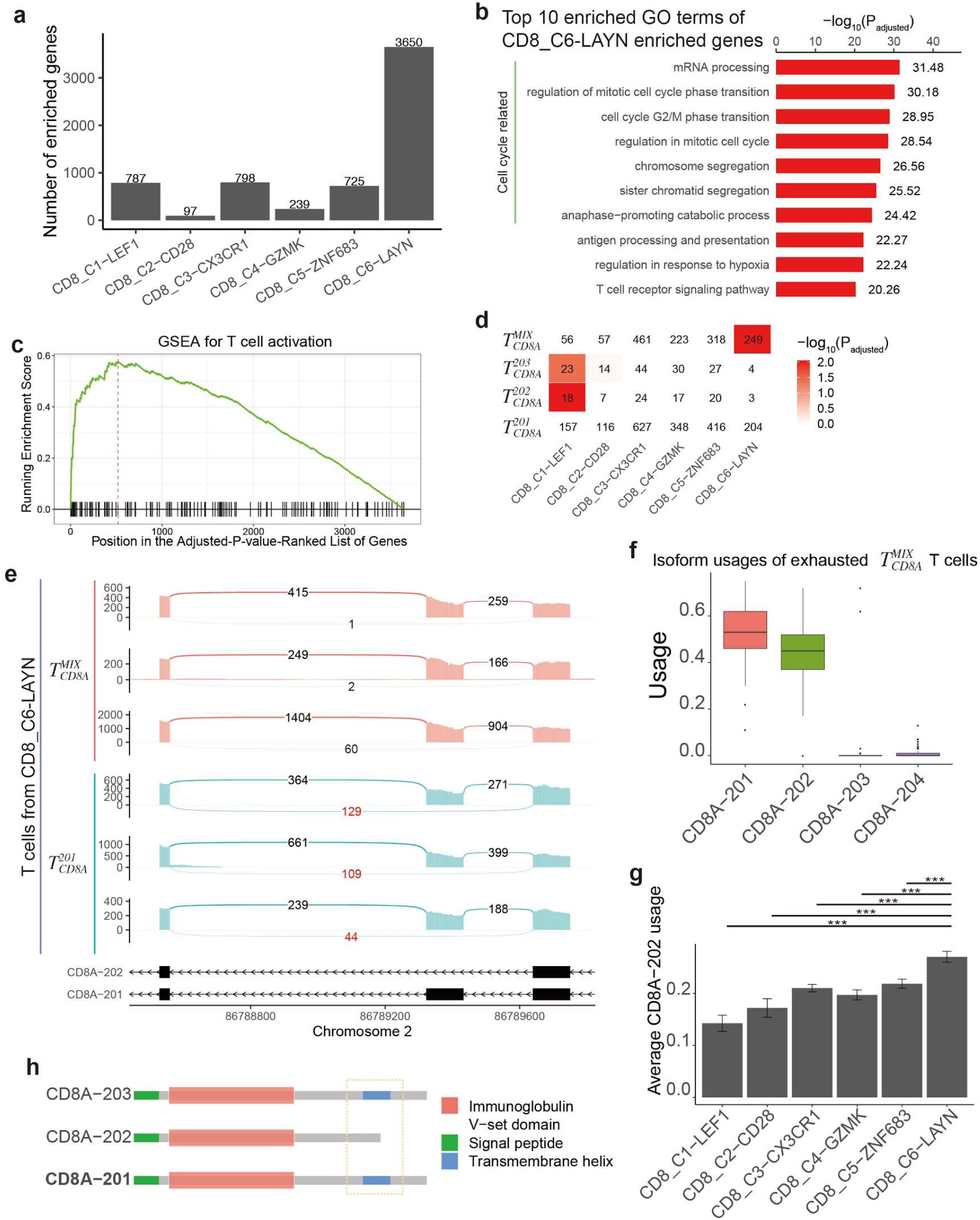
Enrichment of isotypes in CD8+ T cells. **a**, number of enriched genes were counted for each CD8+ T cell subtype. “enriched genes” included any genes with any isotypes detected “enriched” in the given subtypes by enrichment analysis. **b**, bar plots of Gene ontology (GO) terms annotating the enriched genes in CD8 exhausted T cells (CD8_C6-LAYN). Values along the x-axis represent the −log_10_ transformed adjusted p-values, hypergeometric test. **c**, Distribution of running enrichment score of T cell activation pathway highlighted by Gene set enrichment analysis (GSEA). **d**, (lower panel) enrichment results of *CD8A* isotypes. Numbers represent the cell counts shared by each isotype and T cell subtype and colored by −log10 transformed q values from the enrichment analysis. (upper panel) the corresponding protein structures of each enriched isoform. Colored legend indicates protein domains. Dashed yellow box highlights the missing transmembrane helix in CD8A-202 protein isoform. **e**, sashimi plots showing the expression pattern of *CD8A* in CD8A-201 dominated and CD8A-MIX cells. Cells are randomly picked from CD8_C6-LAYN. The numbers on the arcs record the number of junction reads. The red box and numbers highlight reads mapping to exon 2 and the reads skipping it, which is a key feature to distinguish isoform CD8A-202 and CD8A-201. **f**, the distribution of isoform usages in CD8A-MIX exhausted CD8+ T cells. **g**, usage level for CD8A-202 in each CD8+ T cell subtypes. Bar plots showing mean usage level of CD8A-202 in each CD8+ T cell subtypes. By comparing to CD8 exhausted CD8+ T cells, the usage level of other T cell subtypes is significantly lower. Student’s t-test (two-tailed and unpaired) is used to evaluate the statistical significance. **h**, Schematic for protein structure of CD8A protein isoforms. Dashed box indicates the distinct domain among the isoforms.

*CD8A*, a gene encoding a critical co-receptor for T cell receptor, showed differential isoform enrichment in naïve (CD8_C1-LEF1) and exhausted T cells (CD8_C6-LAYN). CD8 specifically binds to class I MHC protein and enhances responses of antigen-specific lymphocytes(32). While most T cells have the isotype of CD8A-201, CD8A-202 and CD8A-203 were significantly enriched in naive CD8+ T cells (CD8_C1-LEF1). In contrast, exhausted CD8+ T cells (CD8_C6-LAYN) were enriched by the T_CD8A_^MIX^ (**Figure 3d**). In almost all of the exhausted T cells with the isotype CD8A-MIX (988/996), CD8A-201 and CD8A-202 were the top two dominant isoforms (**Supplementary Figure 2d**), with the expression level of CD8A-201 slightly higher than CD8A-202 (**Figure 3f**), which is confirmed by higher number of reads supporting CD8A-202 (**Figure 3e**). The enrichment of cells with the T_CD8A_^MIX^ in exhausted CD8+ T cells resulted in high average usage level of CD8A-202 compared with other CD8 T cell subtypes. (**Figure 3g**).

CD8A-202 encodes a protein without the transmembrane region that is present in the canonical isoform CD8A-201 due to the loss of exon 4 (**Figure 3h**, **Supplementary Figure 2g**), indicating that it may be secreted into the extracellular matrix to attenuate the regular CD8+ T cell function. A soluble CD8 isoform excluding exon 4 has been reported in patients with rheumatic disease(33), in patients with EBV-induced infectious mononucleosis(34), and in human cell lines(35). Moreover, the correlation between secretion of CD8 isoform and advanced stages of human diseases such as HIV infection(36), systemic lupus erythematosus(37), rheumatoid arthritis(38), and leukemia(37, 39) has also been reported. Biochemical experiments further proved that soluble CD8 can inhibit CD8+ T cell proliferation and IFN-γ production, attenuate CD8+ T cell responses(40) and impair CD8+ T cell activation and function(41). These findings suggest that soluble *CD8A* in the tumor microenvironment of NSCLC might have the same functions to inhibit CD8+ T cells.

### Functional consequences of differential isotypes between T cell subtypes

To illustrate the functional consequences of differential isotypes between T cell subtypes, we focus on genes with multiple isoforms for the subsequent analysis. A total of 245 and 345 such genes were found for CD8+ and CD4+ T cells, respectively (**Figure 4a and Figure 6a, Supplementary Table 1 and 7**). For CD8+ T cells, exhausted T cells (CD8_C6-LAYN) showed the most distinctions from other T cell subtypes, with 196 genes having significantly differential isotype usage, of which 86 genes were contributed by the differences from tissue resident memory CD8+ T cells (CD8_C5-ZNF683) (**Figure 4b**). Given two T cell subtypes, we compared the gene and protein structures to infer the functional consequences of differential isoform usage. Various databases and tools including ENSEMBL(24), InterPro(25), Uniprot(42), and Pfam(43) were adapted to obtain the functional features of these isoforms, including transcription start site (TSS), coding/non-coding potentials of transcripts, signal peptides, transmembrane region, and protein domains. Based on the annotated functional features, isotype pairs with differential enrichment in T cell subtypes were categorized into five groups with different functional changes including coding/non-coding shifts, alternative TSSs but the same protein sequences, shifts of subcellular locations, and gain/loss of specific domains (**Figure 4a**).

**Figure 4.**
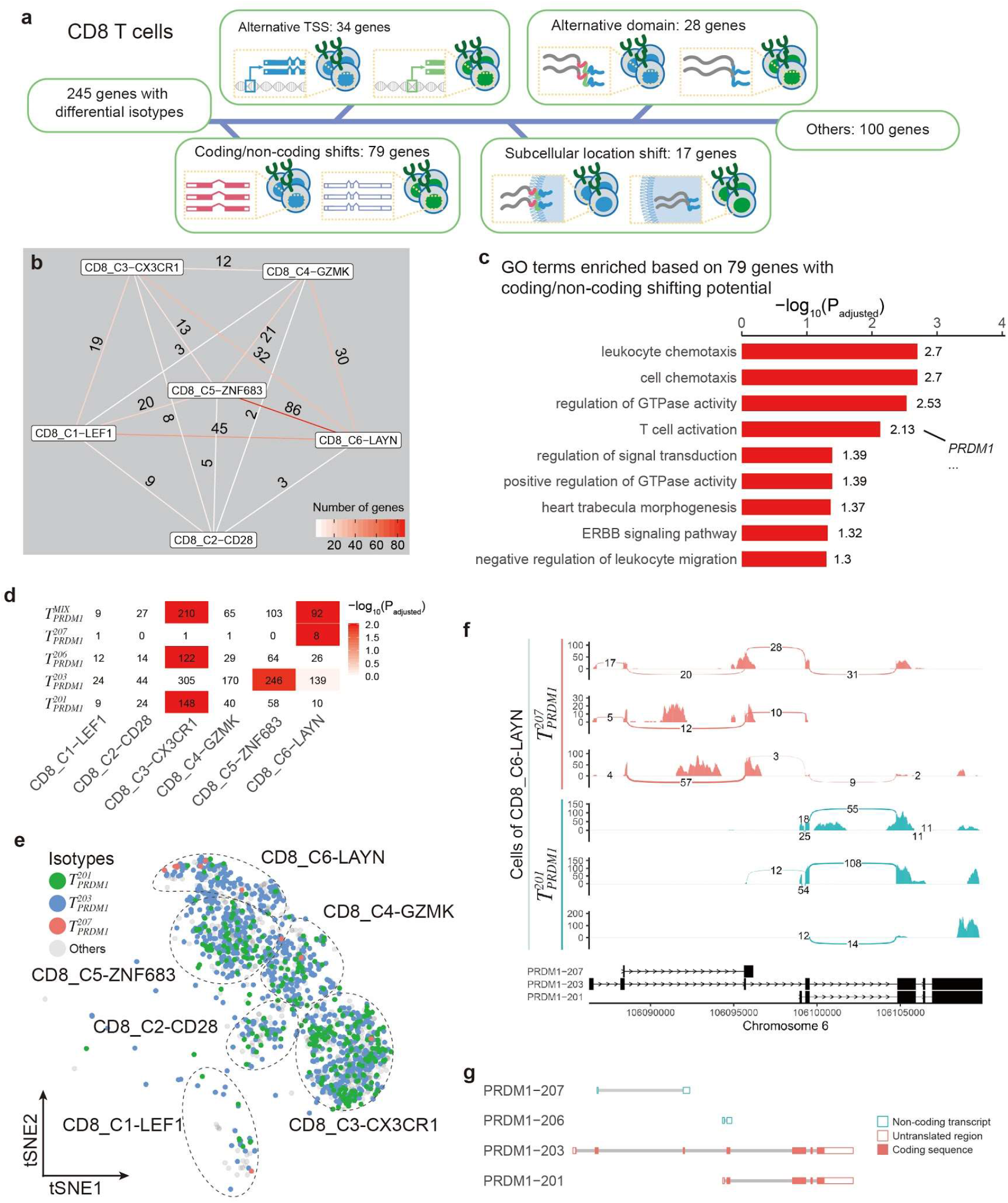
Functional categorizations of differential isotypes in CD8+ T cell subtypes. **a**, Functional categorizations of differential isotypes in CD8+ T cell subtypes based on gene structures. **b**, Numbers of genes (on the edges) with differentially enriched isoforms between CD8+ T cell subtypes. **c**, GO terms enriched by genes within the coding/non-coding shifts category. **d**, Enrichment of *PRDM1* involved in T cell activation between CD8+ T cell subtypes. Numbers: cell counts colored by −log_10_ transformed adjusted p values. **e**, tSNE plot showing the cell isotypes for *PRDM1* in CD8+ T cells. **f**, sashimi plots showing the expression pattern of *PRDM1* in T_PRDM1_^201^ and T_PRDM1_^207^ T cells in CD8_C6-LAYN subtype. Numbers on the arcs the number of junction reads. **g**, Structure of transcripts of *PRDM1* isoforms.

For genes with coding/non-coding shifts (**Supplementary Table 2**), pathways such as chemotaxis and T cell activation were enriched based on gene ontology enrichment analysis (**Figure 4c**), of which *PRDM1* is of particular interest for its crucial roles in CD8+ T cell differentiation and exhaustion(44). The isotypes of effector memory (CD8_C3-CX3CR1) and exhausted T cells (CD8_C6-LAYN) for *PRDM1* showed significant differences (**Figure 4d**), with the effector T cells enriched T_PRDM1_^201^ and T_PRDM1_^206^, and T_PRDM1_^MIX^, while exhausted T cells enriched T_PRDM1_^207^ and T_PRDM1_^MIX^ (**Figure 4d**). The enrichment of T_PRDM1_^207^ cells in the exhausted T cell subtype can also be readily observed in the t-SNE visualization (**Figure 4e**). The pre-exhausted T cell subtype (CD8_C5-ZNF683) showed a preference for T_PRDM1_^203^ cells. Close examination of reads mapping to the gene region of *PRDM1* confirmed the accuracy of the isotype determination exemplified by T_PRDM1_^207^ and T_PRDM1_^201^ in exhausted T cells (**Figure 4f**). Previous study suggested that *PRDM1* can upregulate the expression of PD-1 and TIGIT and thus impair the T cell function(45). *PRDM1* was also known to direct virus-specific effector T cells to differentiate into exhausted instead of memory T cells during chronic lymphocytic choriomeningitis virus infection(46). Our results of the differential isoform usage of *PRDM1* between effector memory and exhausted T cells may suggest that the splicing of *PRDM1* plays important roles in the regulation of tumor-infiltrating T cell exhaustion.

GO term enrichment analysis for those genes belonging to the “alternative TSS” category (**Supplementary Table 3**) suggests that biological processes including T cell migration and protein localization to cell periphery were enriched, indicating the possible regulatory role of splicing in these genes (**Figure 5a**). Among these genes, isoforms of *CCR6*, which is responsible for the chemotaxis of T cells(47), were found to be differentially enriched by CD8_C4-GZMK and CD8_C5-ZNF683 (**Figure 5b**), with CCR6-201 dominantly expressed by CD8_C4-GZMK and CCR6-203 dominantly expressed by CD8_C5-ZNF683 (**Figure 5c**). Both isoforms have the same protein product but different transcription start sites (TSS). While these two T cell subtypes both have less exhausted features, they are distinct from each other regarding the signatures of tissue-resident memory T cells. The preferences for different TSSs in these two subtypes(3) may provide the isoform-level clues for the molecular mechanisms of tissue-resident memory T cells.

**Figure 5.**
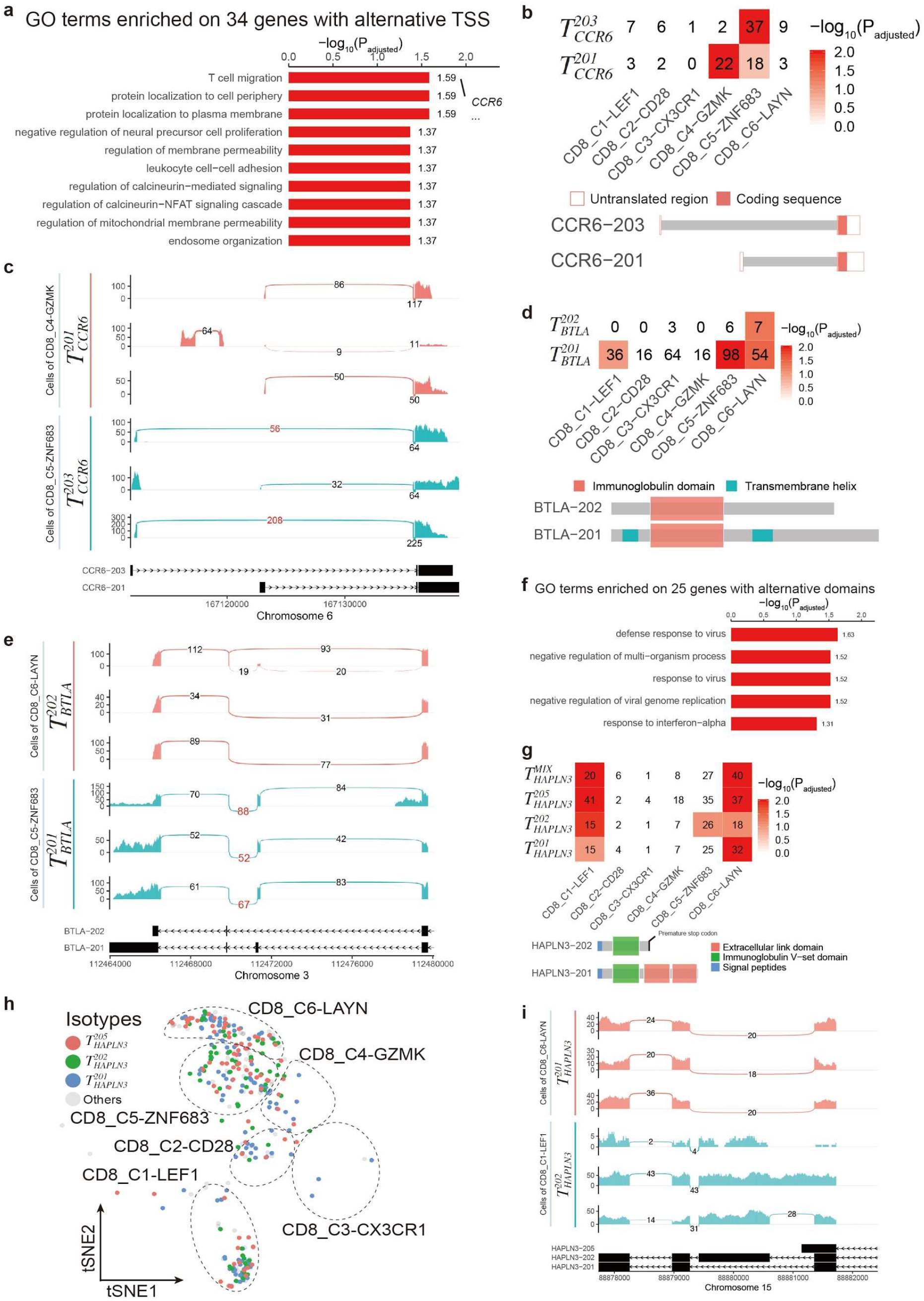
Genes with differential isotypes in other functional categories. **a**, GO terms enriched by genes within alternative TSS category. **b**, Enrichment of CCR6 involved in T cell migration between CD8+ T cell subtypes (the upper panel) and the corresponding transcript structures of each enriched isoform (lower panel). Numbers: cell counts colored by −log10 transformed adjusted p values. **c**, sashimi plots showing the expression pattern of *CCR6* in T_CCR6_^201^ and T_CCR6_^203^ cells. **d**, Enrichment results of *BTLA* (the upper panel) and the corresponding protein structures of each enriched isoform (lower panel). **e**, sashimi plots showing the expression pattern of *BTLA* in T_BTLA_^201^ and T_BTLA_^202^ cells. **f**, GO terms enriched by genes within alternative domain category. **g**, Enrichment results of *HAPLN3* isotypes (the upper panel) and the corresponding protein structures of each enriched isoform (lower panel). **h**, tSNE plot showing the isotypes for *HAPLN3*. **i**, sashimi plots showing the expression pattern of *HAPLN3* in T_HAPLN3_^201^ and T_HAPLN3_^202^ dominated cells. The side bars in sashimi plots denotes the clusters of the selected cells. CD8_C1-LEF1, naive CD8+ T cells; CD8_C4-GZMK, pre-exhausted CD8+ T cells; CD8_C5-ZNF683, tissue resident memory CD8+ T cells; CD8_C6-LAYN, exhausted CD8+ T cells. Numbers on the arcs in sashimi plots denote the number of junction reads. The red numbers highlight the reads mapping to key junctions to distinguish the compared isoforms.

**Figure 6.**
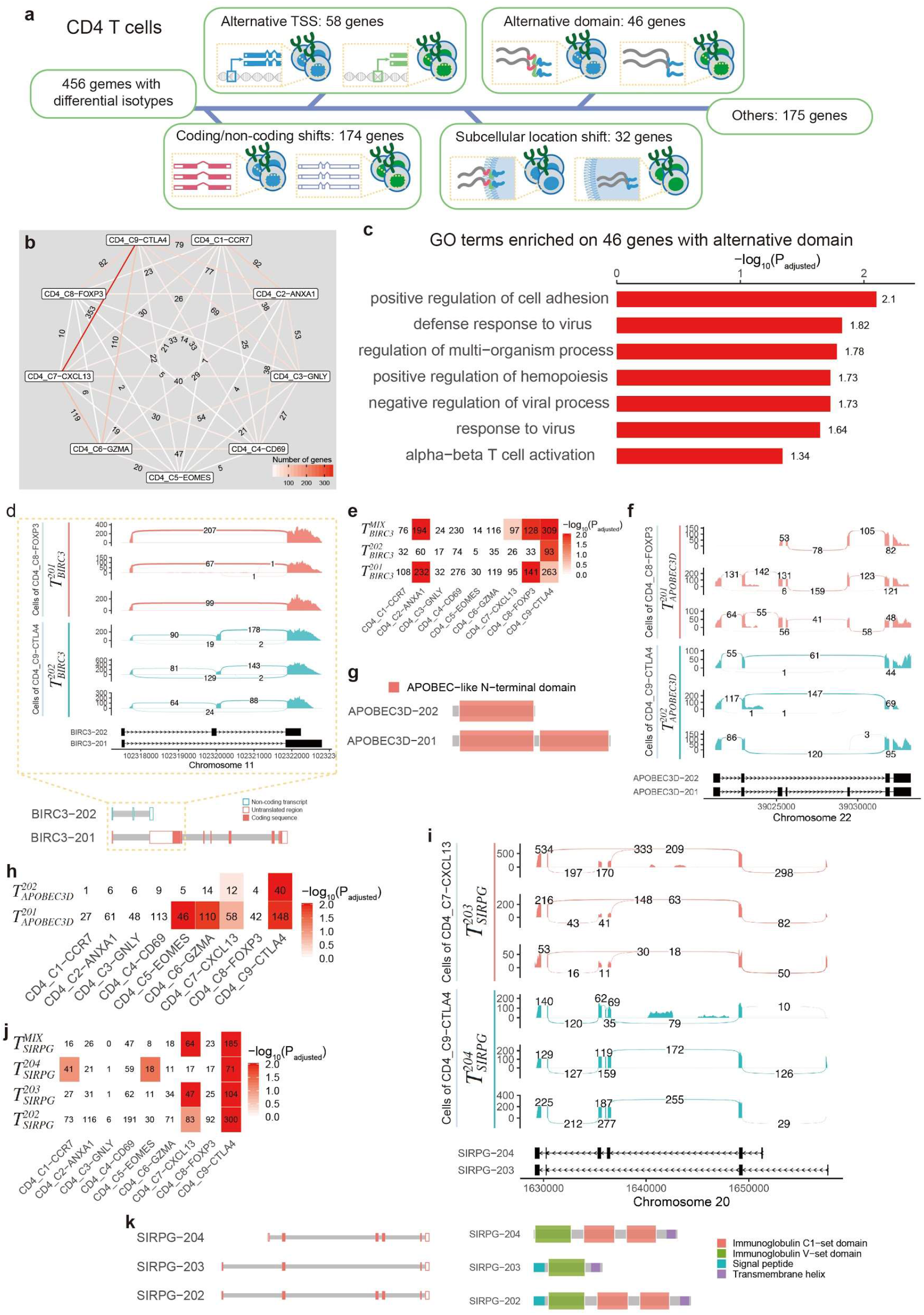
Differentially enriched isotypes in CD4+ T cell subtypes. **a**, Functional categories of differentially enriched isotypes in CD4+ T cell subtypes. **b**, Numbers of genes with differentially enriched isotypes between two CD4+ T cell subtypes. **c**, GO terms enriched by the genes within alternative domains category. **d**, sashimi plots showing the splicing patterns of *BIRC3* in regulatory T cells (the upper panel) and transcript structures of each enriched *BIRC3* isoforms in **e** (lower panel). **e**, enrichment of *BIRC3* isotypes in CD4+ T cells. Numbers represent the cell counts in each category and colored by −log_10_ (P_adjusted_) from the enrichment analysis. **f**, sashimi plots showing the splicing pattern of *APOBEC3D* in regulatory T cells. **g**, the protein domain structures of each enriched *APOBEC3D* isoform in **h**. **h**, enrichment of *APOBEC3D* isotypes in CD4+ T cells. **i**, sashimi plots showing the splicing pattern of *SIRPG* in regulatory and helper T cells. **j**, enrichment of *SIRPG* isotypes. **k**, the protein domain structures of each enriched *SIRPG* isoform in **j**. The numbers on the arcs in sashimi plots record the number of junction reads. The red numbers and boxes highlight the reads mapping to key junctions and exons respectively.

Isoforms of 17 genes with differential preferences in CD8+ T cells were detected to have differed subcellular location (**Supplementary Table 4**). For example, *BTLA*, which negatively regulates antigen receptor signaling(48, 49) and helps naive T cells to remain in resting state(50), was observed to have two significantly enriched isoforms differed by a transmembrane domain. The canonical isoform BTLA-201, encoding a membrane-integrated protein, was observed to be dominantly expressed by T cells of CD8_C1-LEF1, CD8_C5-ZNF683, and CD8_C6-LAYN (**Figure 5d**). In addition to BTLA-201, a significant proportion of exhausted T cells (CD8_C6-LAYN) dominantly expressed BTLA-202.The dominantly expressed BTLA-202 in T_BTLA_^202^ cells were confirmed by examination of the reads coverage by sashimi plots (**Figure 5e**), with exon 3 excluded during splicing. The splicing of BTLA-202 leads to loss of the transmembrane helix domain, resulting in a soluble *BTLA* product compared with the canonical isoform. This soluble isoform was confirmed and found to increase in the early stages of sepsis(51, 52) and was able to enhance antitumor activity in a melanoma pulmonary metastasis model in combination of HSP70 vaccine(53). Thus, the enrichment of T_BTLA_^202^ cells in CD8+ exhausted T cells may suggest a new mechanism for the induction of immunosuppressive tumor microenvironment.

25 genes that have alternative domains between isoforms were found with differential enrichment among CD8+ T cell subtypes (**Supplementary Table 5**). GO term enrichment analysis suggested that these genes participate in immune responses to viral infection (**Figure 5f**).

One representative gene is *HAPLN3*, which was highly expressed in naive T cells (CD8_C1-LEF1) and exhausted T cells (CD8_C6-LAYN) (**Supplementary Figure 1d**). In both T cell subtypes, multiple isotypes were observed, including T_HAPLN3_^201^, T_HAPLN3_^202^, T_HAPLN3_^205^, and T_HAPLN3_^MIX^. While T_HAPLN3_^205^ and T_HAPLN3_^MIX^ were enriched in both of the two T cell subtypes, naive T cells were more enriched by T_HAPLN3_^202^ cells and exhausted T cells were more enriched by T_HAPLN3_^201^ cells (**Figure 5g,h**). Compared with HAPLN3-201, HAPLN3-202 has an extra exon 4 (**Figure 5i**), which results in a premature stop codon and consequently nonsense-mediated decay of the transcript(24). The protein product of HAPLN3-202 loses the extracellular link domain which inhibits the cell adhesion function. These differences between naive T cells and exhausted T cells regarding the isoforms of HAPLN3 may be associated with the migratory and tumor-resident properties of naïve and exhausted T cells, respectively, although the expression levels were similar at the gene level.

Analyses at isoform level also provide new insights into gene functions in CD4+ T cell subtypes. A total of 442 genes were detected with differential isotypes between CD4+ T cell subtypes (**Figure 6a**), including 177 genes with coding/non-coding differences between isoforms (**Supplementary Table 8**), 58 genes with alternative TSSs (**Supplementary Table 9**), 32 genes with distinct subcellular location potentials (**Supplementary Table 10**), and 46 genes with alternative domains (**Supplementary Table 11**). Across all the 9 CD4+ T cell subtypes, we found that tumor-infiltrating regulatory T cells (CD4_C9-CTLA4) have the most distinct pattern at isoform level compared with other subtypes and 353 genes were detected with differential isotypes between regulatory T cells and exhausted CD4+ T cells (CD4_C7-CXCL13) (**Figure 6b**). Biological pathways involved in responses to viral infection were enriched in genes with alternative domains between isoforms detected in CD4+ T cells (**Figure 6c**).

*BIRC3* is one of the 177 genes that have isoforms with different coding potentials (**Supplementary Table 8**), with BIRC3-202 having no open reading frames within the transcript according to the ENSEMBL annotation. Reads mapping confirmed the existence of its second exon unique in BIRC3-202 and supported that the isotypes of these cells were T_BIRC3_^202^ (**Figure 5d**). This isotype was enriched by tumor-infiltrating regulatory T cells (CD4_C9-CTLA4) while central memory CD4+ T cells (CD4_C2-ANXA1) and resting regulatory CD4+ T cells from blood (CD4_C8-FOXP3) enriched the isotype T_BIRC3_^201^ (**Figure 5e**). The coding/non-coding differences in blood- and tumor-derived regulatory T cells may suggest *BIRC3* plays an important role in shaping the tumor immune microenvironment.

*APOBEC3D* is a gene encoding a DNA deaminase which acts as an inhibitor of retrovirus replication and retrotransposon mobility(54, 55). Our study revealed that the isoforms of *APOBEC3D* have different preference in CD4+ T cells, with tumor-infiltrating regulatory T cells (CD4_C9-CTLA4) significantly enriched T_APOBEC3D_^202^ cells in addition to T_APOBEC3D_^201^ compared with other CD4+ T cell subtypes (**Figure 6h**). Different from APOBEC3D-201, APOBEC3D-202 loses the entire APOBEC-like domain by skipping exon 3, 4 and 5, which is confirmed by sashimi plots (**Figure 6f, g**). Such enrichment of T_APOBEC3D_^202^ cells in tumor-infiltrating CD4+ T cells including Tfh-like T cells (CD4_C7-CXCL13, not significant) and regulatory T cells (CD4_C9-CTLA4) may suggest that tumor microenvironment can change the splicing scenarios of the infiltrating T cells but further exploration is needed to clarify this hypothesis.

*SIRPG* can interact with CD47 expressed by antigen-presenting cells and regulate T cell activation and proliferation(56). Compared with other CD4+ T cell subtypes, tumor-infiltrating regulatory T cells (CD4_C9-CTLA4) enriched T_SIRPG_^202^, T_SIRPG_^203^, T_SIRPG_^204^, and T_SIRPG_^MIX^ cells, and Tfh-like cells (CD4_C7-CXCL13) enriched T_SIRPG_^202^, T_SIRPG_^203^, and T_SIRPG_^MIX^ cells (**Figure 6j**). These three isoforms were all detected with transmembrane helix but differed by the signal peptide and immunoglobulin C1-set domains (**Figure 6k**). These differences are caused by varied usage of exon 3 and 4, which is also confirmed by reads mapped to the critical junctions and exons (**Figure 6i**). The various isotypes of tumor-infiltrating CD4+ T cells for *SIRPG* suggest the great heterogeneity of tumor-infiltrating T cells at the transcript isoform level with single-cell resolution and probably provide new clues of immunoediting in the tumor microenvironment of lung cancers which needs further exploration.

### Pervasive isoform switching across multi-isoform genes

Isoform switching between conditions was frequently reported by previous studies based on bulk samples(57). However, evidence at the single-cell resolution has been rarely reported. Here we utilized the clonal information of T cells informed by TCR sequences to identify isoform switching events on the single-cell level. Since T cells with the same TCR sequences were originated from the same clone, given one gene, the occurrence of more than one isotype within the same T cell clone would indicate isoform switching.

A total of 3020 T cells were found to share identical TCRs with other cells, including 1988 CD8+ and 1032 CD4+ clonal T cells (**Figure 7a**). CD8+ clonal T cells formed 379 TCR clonotypes with the number of clonal cells within each clonotype ranging from 2 to 66. 280 clonotypes were identified within CD4+ clonal T cells and the number of cells within each clonotypes ranges from 2 to 72. For these clonotypes, we found that a large proportion of genes have more than two isotypes in a single clone (**Figure 7b**). Among these genes, *FYN* was found to be the most diverse genes, with 13 isotypes identified in clone P0616A_C000008:88. Among the 583 clonotypes that had FYN expressing, 26% (153/583) of clones were found to have at least two isotypes for *FYN* (**Figure 7c**). Beyond *FYN*, 10032 out of 14629 genes with multiple isoforms expressed in CD8+ clonal T cells were also found to have at least two isotypes in at least one clonotype (**Figure 7d**). Similar observations were also found in CD4+ T cell clonotypes, with 64% (9328/14629) of genes having at least two isotypes identified in at least one clonotype. Together, these observations indicate that pervasive isoform switching events occurred during T cell activation/proliferation in the lung cancer tumor microenvironment, thus the genes having at least two isotypes in at least one clonotype were termed as genes with isoform switching. Close examination of *CD8A* and *APOBEC3D*, two of genes with isoform switching, in individual cells confirmed the isoform switching within the same T cell clonotypes (**Supplementary Figure 3**). In the clonotype P0616A_C000008:88 with 88 T cells, four CD8A isotypes were found and each was supported by abundant reads mapped to the specific junctions (**Supplementary Figure 3a, b**). Similarly, three isotypes of *APOBEC3D* were robustly identified in CD4+ T cell clonotypes exemplified by P1202_C000002:95 (**Supplementary Figure 3c, d**). In addition, the genes with differential enriched isotypes were extremely enriched in genes with isoform switching in both CD8 T cells and CD4 T cells (**Figure 6f, g**). Altogether, these results suggest that isoform switching is a pervasive phenomenon during T cell clonal expansion and may contribute to the differential isoform usage in various T cell subtypes that were formerly observed.

**Figure 7.**
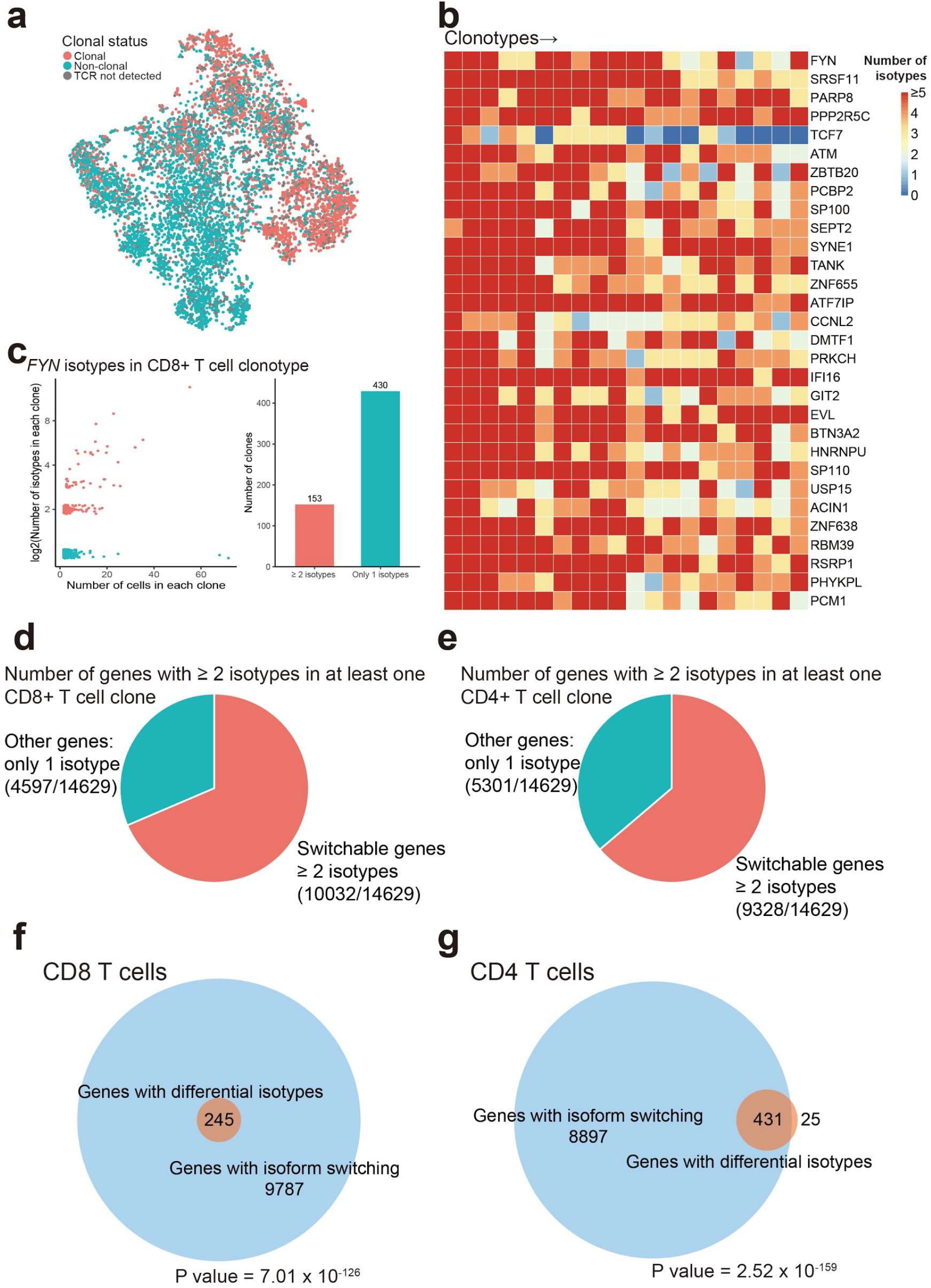
Isoform switching observed in multi-isoform genes based on T cell clones. **a**, Clonal status of both CD8+ and CD4+ T cells showing on tSNE space. **b**, The number of isotypes found for top 30 genes with most expressed isotypes in each T cell clonotype. Each column corresponds to a TCR clone (Clone name not shown). **c**, Number of *FYN* isotypes found in each CD8+ T cell clonotype. Small amount of variation was added to the points along the y-axis. The bar plot on the left showing the corresponding numbers of genes in each category. The proportion of genes with multiple isotypes in at least two CD8+ T cell clones (**d**) or two CD4+ T cell clones (**e**). Venn diagram of genes with differential enriched isotypes and genes with isoform switching in (**f**) CD8+ T cells and (**g**) CD4+ T cells. Numbers indicate the number of genes. Hypergeometric test is used to evaluate the statistical significance.

## DISCUSSION

In this study, we proposed a computational pipeline to investigate the isoform landscape at single-cell resolution, applied it to explore the expression landscape of tumor-infiltrating T cells of non-small cell lung cancer patients, and made several important findings. First, in single T cells, genes with multiple transcript isoforms were demonstrated to dominantly express one isoform compared with bulk RNA-seq data. This finding is consistent with previous observations based on single cells(28, 29), suggesting significant biological differences observed in single cells in contrast to observations based on bulk studies. The wide existence of dominantly expressed isoforms in single cells also highlights the great phenotypic heterogeneity of cells, providing a new dimension to investigate the genotype-phenotype relationship, particularly in the tumor context.

Given a gene, the wide existence of dominantly expressed isoforms in single cells allows defining the isotypes of a cell to statistically investigate the isoform preferences among T cell subtypes. Based on Fisher’s exact test, we established a statistical pipeline to quantify the significance of isoform preference between T cell subtypes and observed that 557 genes expressed differential isoforms among CD8+ T cell subtypes with sufficient significance after multiple testing correction. Even for genes constitutively expressed in CD8+ T cells such as *CD8A*, significant changes at the isoform level were also detected among T cell subtypes, suggesting that differential isoform usage may be a widely applied mechanism to regulate the function and state of T cells. In particular, most differential isoform usage events were found between the tumor-infiltrating exhausted CD8+ T cells and other CD8+ T cells and between the tumor-infiltrating CD4+ Tregs and other CD4+ T cells, which may suggest that changing the alternative splicing landscape is another important mechanism for tumor microenvironment co-opting the infiltrated T cells.

Functional annotations of the isotypes differentially enriched in different T cell subtypes revealed that a set of diverse biological processes were impacted by the differential isoform usage, including coding/non-coding switches, alternative TSS, protein domain gains/losses, and subcellular translocations. Some of the critical genes such as *CD8A*, *PRDM1*, *CCR6*, *BTLA* and *HAPLN3* that have been previously related to T cell dysfunction via alternative splicing were observed to have distinct isotypes in tumor-infiltrating exhausted T cells compared with other T cell subtypes and were involved in different biological processes, indicating the high heterogeneity of tumor immune microenvironment. A remarkable example among these genes is *CD8A*, of which its soluble isoform (CD8A-202) was previously found to be expressed by CD8+ T cells during several human diseases such as HIV infection(36), systemic lupus erythematosus(37), rheumatoid arthritis(38), and leukemia(37, 39), has the potential to compete with membrane-integrated *CD8A* protein and thus impairs the cytotoxicity of CD8+ T cells(40, 41). The enrichment of exhausted CD8+ T cells with the soluble CD8A as the isotype in the non-small cell lung cancer may enable the immune suppression status of the tumor microenvironment. Further experimental explorations are needed to validate this tempting hypothesis.

We further employed the clonal information of T cells derived from the sequences of T cell receptors to investigate whether the differential isotypes among T cell subsets were developmentally independent or derived from isoform switching during T cell activation/differentiation. The results suggested that pervasive genes with multiple isoforms expressed in T cells underwent isoform switching during T cell clonal expansion, including almost all those genes showing differential isotypes among T cell subtypes. These observations suggest that isoform switching is a common phenomenon during T cell activation/differentiation in the tumor microenvironment. It is necessary to consider the isoform information to explain the efficacy of the current immunotherapies and to develop future immunotherapies that target to tumor-infiltrating T cells.

Overall, we established an analytical framework to investigate the isoform landscape of tumor-infiltrating T cells at the single-cell resolution and revealed the existence of isotypes for genes with multiple isoforms in individual cells. The extensive isoform switching during T cell clonal expansion and differential isotype usage among T cell subsets involving a set of diverse biological processes are reminders of the great phenotypic heterogeneity of tumor-infiltrating T cells. Furthermore, this vast heterogeneity drives the necessity of single-cell isoform analytics for understanding tumor T cell immunity. The findings uncovering various isoform switching events provide a rich data resource for deep exploration of the tumor immune microenvironment.

Finally, there is no reason to believe that the differential isoform usage and switching is restricted to T cells. We identified T cell isoform patterns because we were at the advantageous position to have generated such deeply sequenced single cell SMART-seq2 data. Despite the wide prevalence of scRNA-seq data, most datasets (e.g. those by the 10x Genomics and other drop-seq platforms) might not have the depth to reliably reveal splicing differences. Those who take on the strenuous challenge to generate the deep SMART-seq2 and other longer-read data should be encouraged to examine the complex and important isoform usage and switching landscape in other biological systems.

## DATA AVAILABILITY

The data used in this study can be accessed from European Genome-phenome Archive database (accession number EGAS00001002430).

## ACKNOWLEDGMENTS

We thank Hannah Comeau for critical reading of the manuscript. We also thank members of Zhang laboratory for helpful comments and discussions. We thank the Computing Platform of the CLS (Peking University) for providing computing resource.

---

**Supplementary Figure 1.**
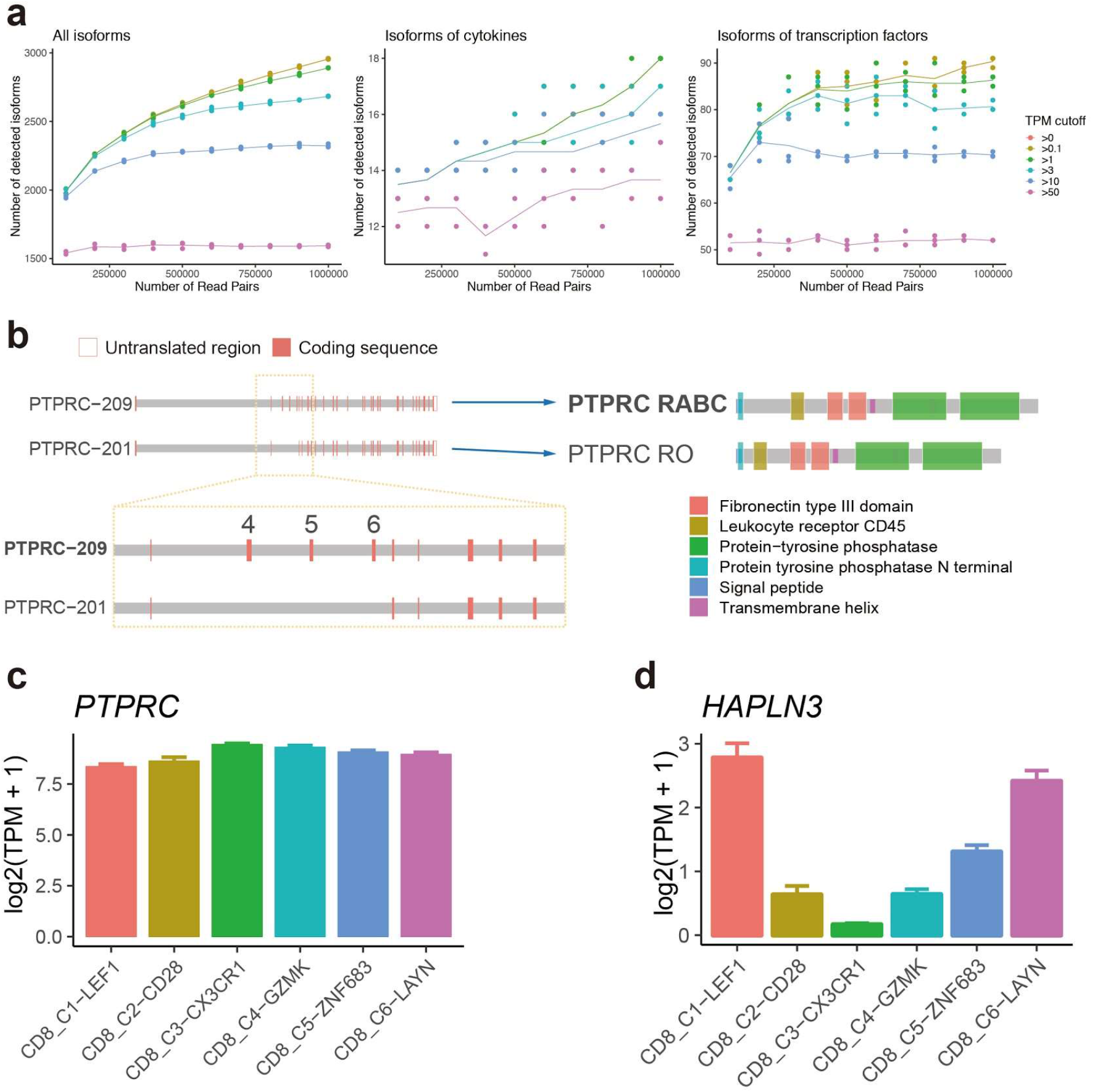
**a**, Saturation curves for the number of isoforms detected at different expression levels. Each point on the curve is derived from calculations based on the random selection of a fraction of raw reads from each sample. Each line with different color shows how fast a isoform can reach detection saturation at different expression levels, represented by a particular TPM value. For isoforms with TPM > 3, 500000 reads are sufficient for detecting the vast majority of such genes. For low expressing isoforms of genes such as cytokines and transcription factor, the curves are more vulnerable to fluctuation. N = 3. **b**, Structure of transcripts of *PTPRC* isoforms and the corresponding protein products of each transcript. Gene expression of *PTPRC* (**c**) and *HAPLN3* (**d**) were shown among CD8 T cell subtypes. Error is s.e.m.

**Supplementary Figure 2.**
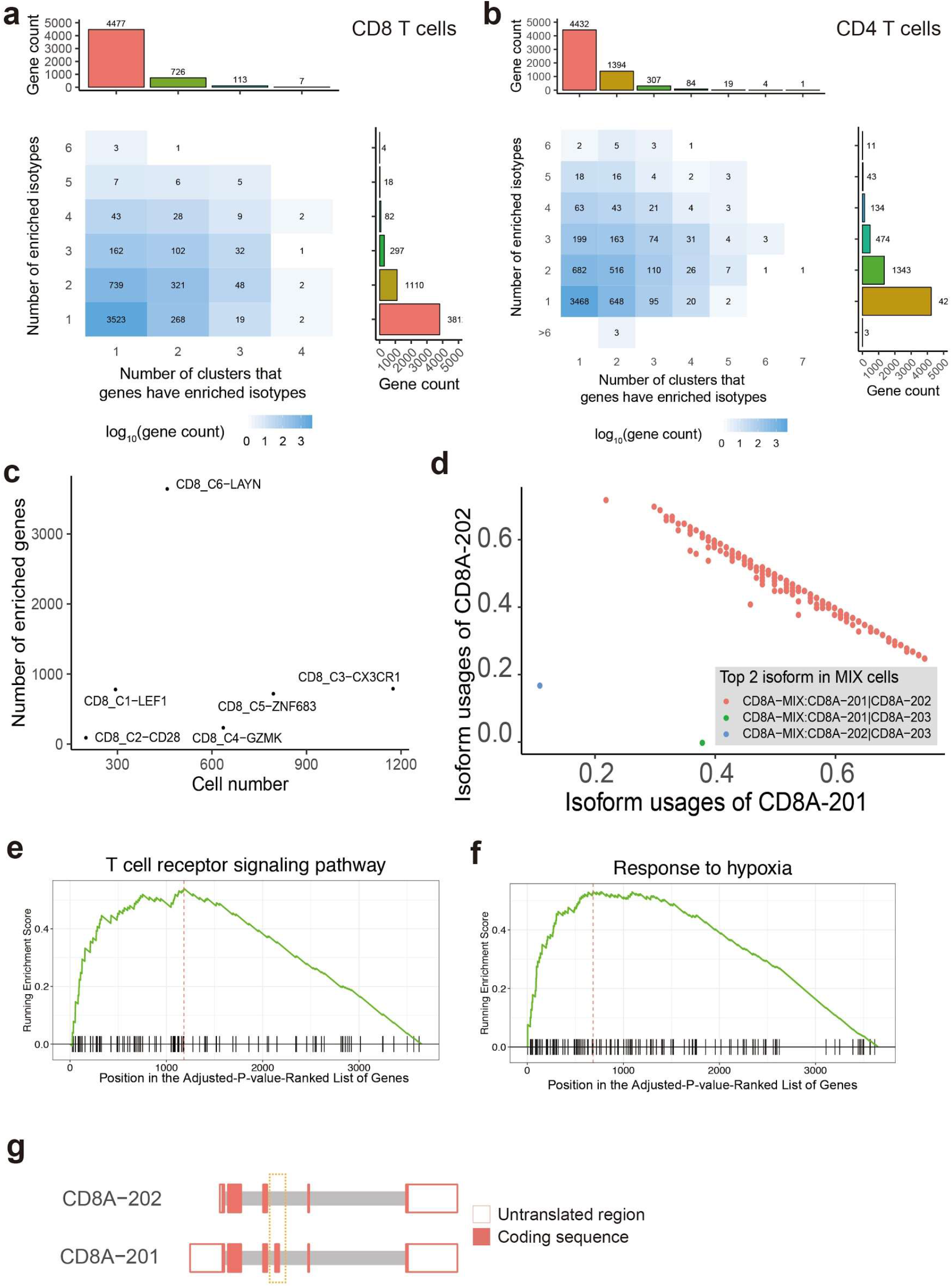
Number of enriched genes in CD8+ (**a**) and CD4 T cell subtypes (**b**). **c**, dot plot showing no obvious association between number of enriched genes and number of cells in each CD8 T cell subtype. **d**, Most of CD8A-MIX cells are expressing CD8A-201 and CD8A-202 simultaneously. Each dot denotes a T_CD8A_^MIX^ cell colored by the top 2 highly expressed CD8A isoform in it. Distribution of running enrichment score of T cell receptor signaling pathway (**e**) and response to hypoxia (**f**) highlighted by Gene set enrichment analysis (GSEA). **g**, Structure of transcripts of *CD8A* isoforms. Dashed box indicates the distinction between two isoforms.

**Supplementary figure 3.**
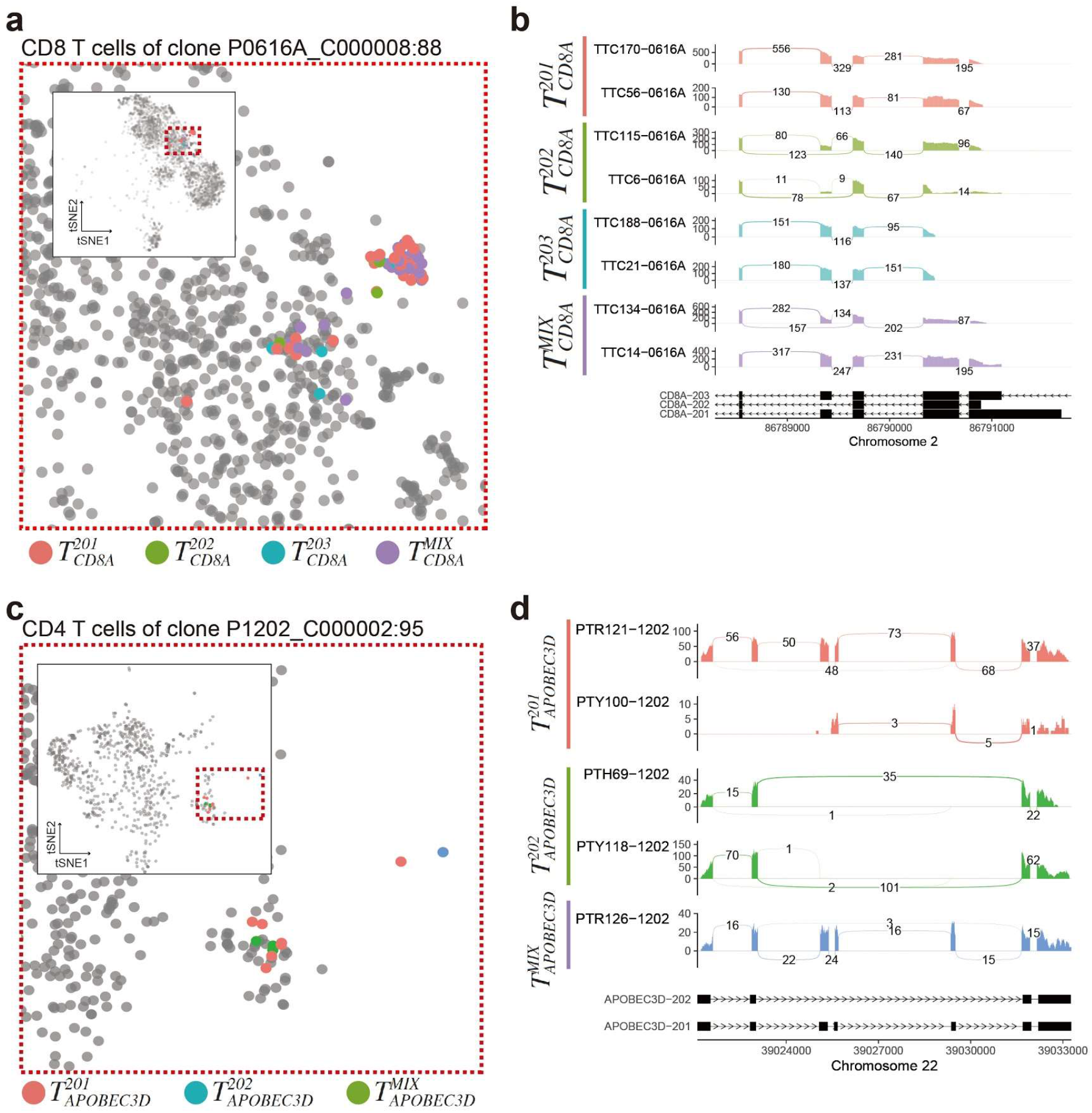
**a**, Clones with multiple *CD8A* isotypes are exemplified by clone P0616A_C000008:88. T cells belong to this clone are highlighted in tSNE plot and colored by isotypes. **b**, Sashimi plot showing the distinctive expression pattern of *CD8A* isotypes of cells in clone P0616A_C000008:88. **c**, Clones with multiple *ABOPEC3D* isotypes are exemplified by clone P1202_C000002:95. **d**, Sashimi plot showing the distinctive expression pattern of *APOBEC3D* isotypes of cells in clone P1202_C000002:95.

