## Supplementary figures and images for "Landscape of transcript isoforms in single T cells infiltrating in non-small cell lung cancer"

### Supplemental Figure 1

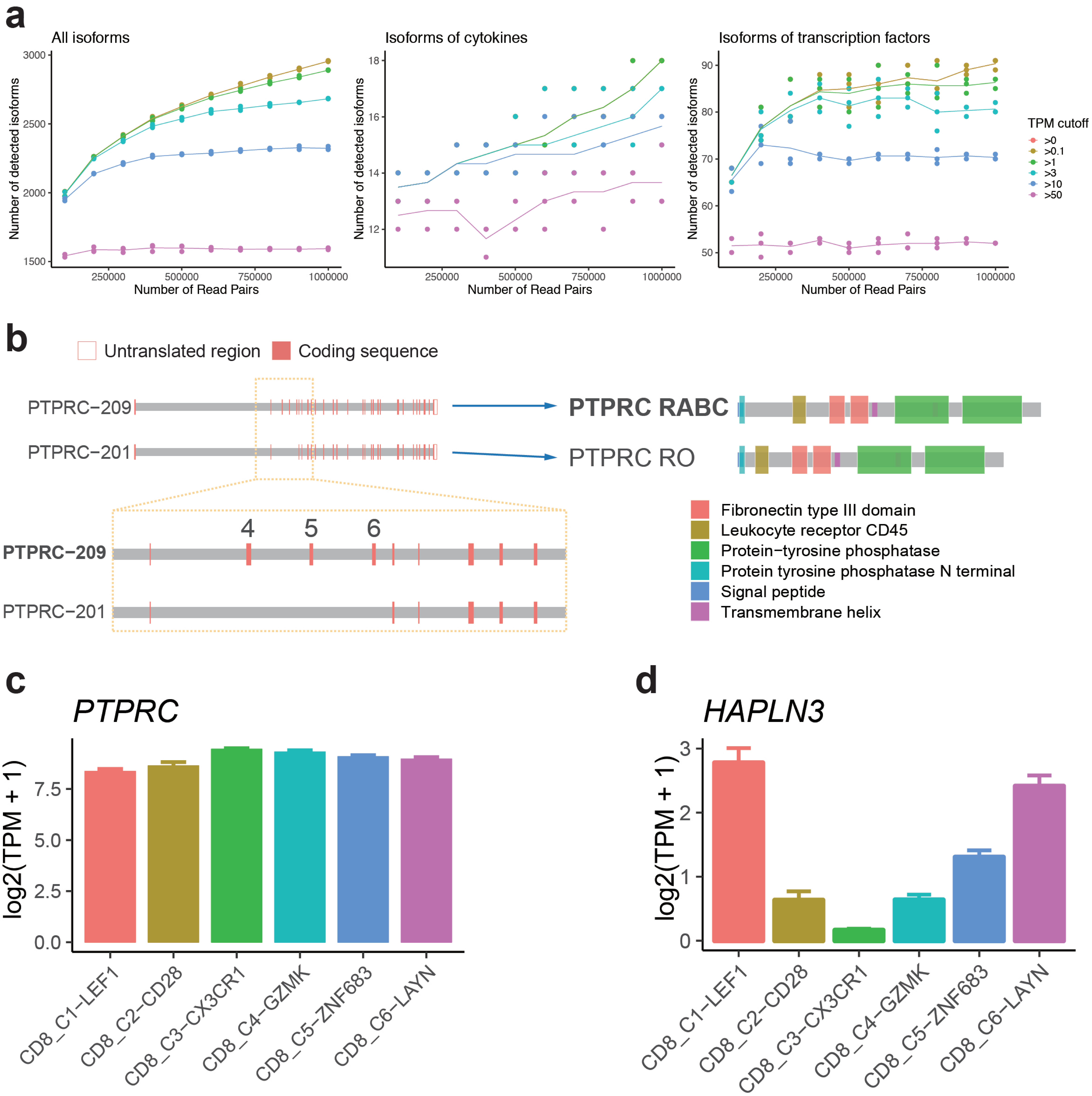

### Supplemental Figure 2

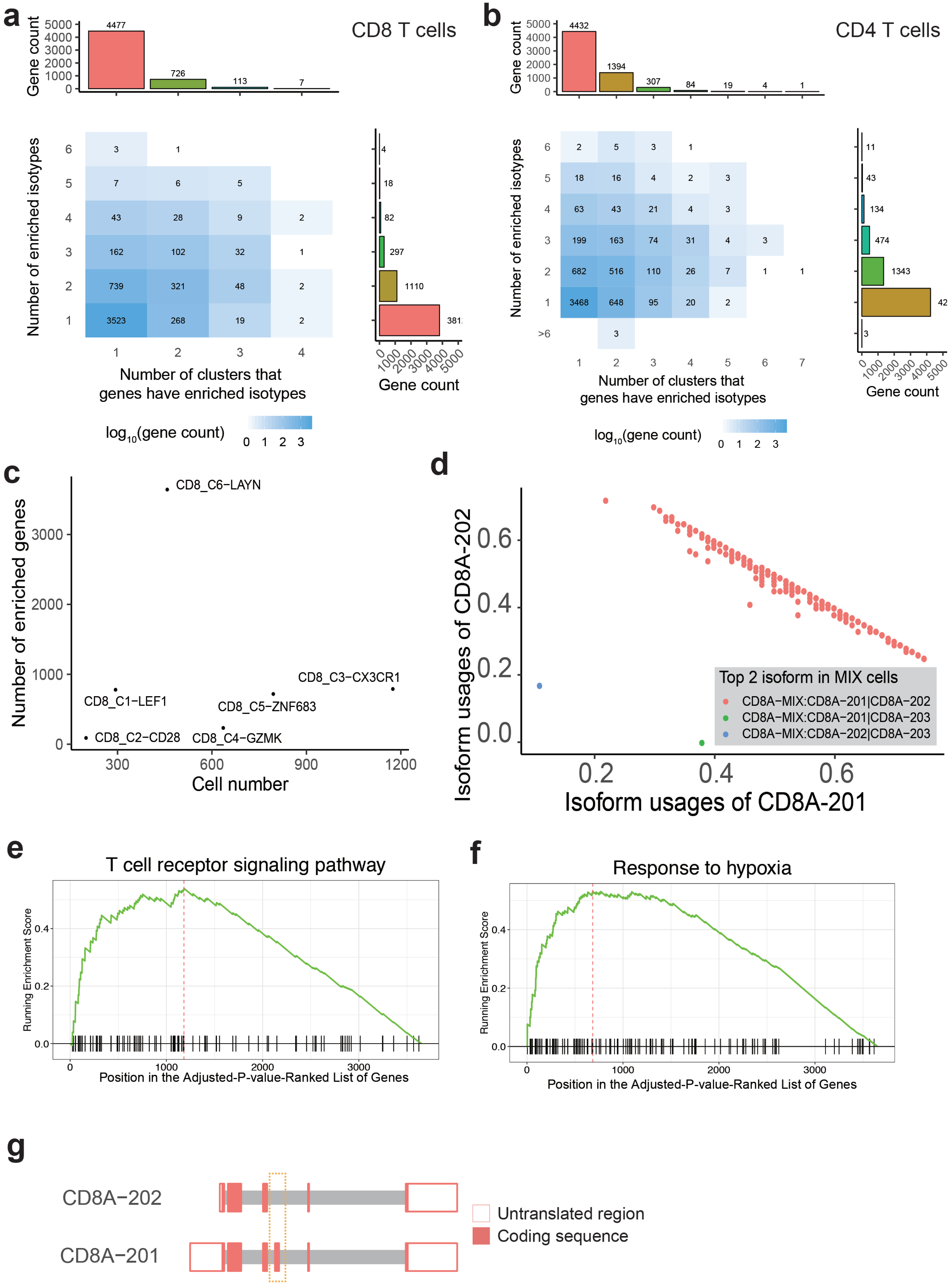

### Supplemental Figure 3

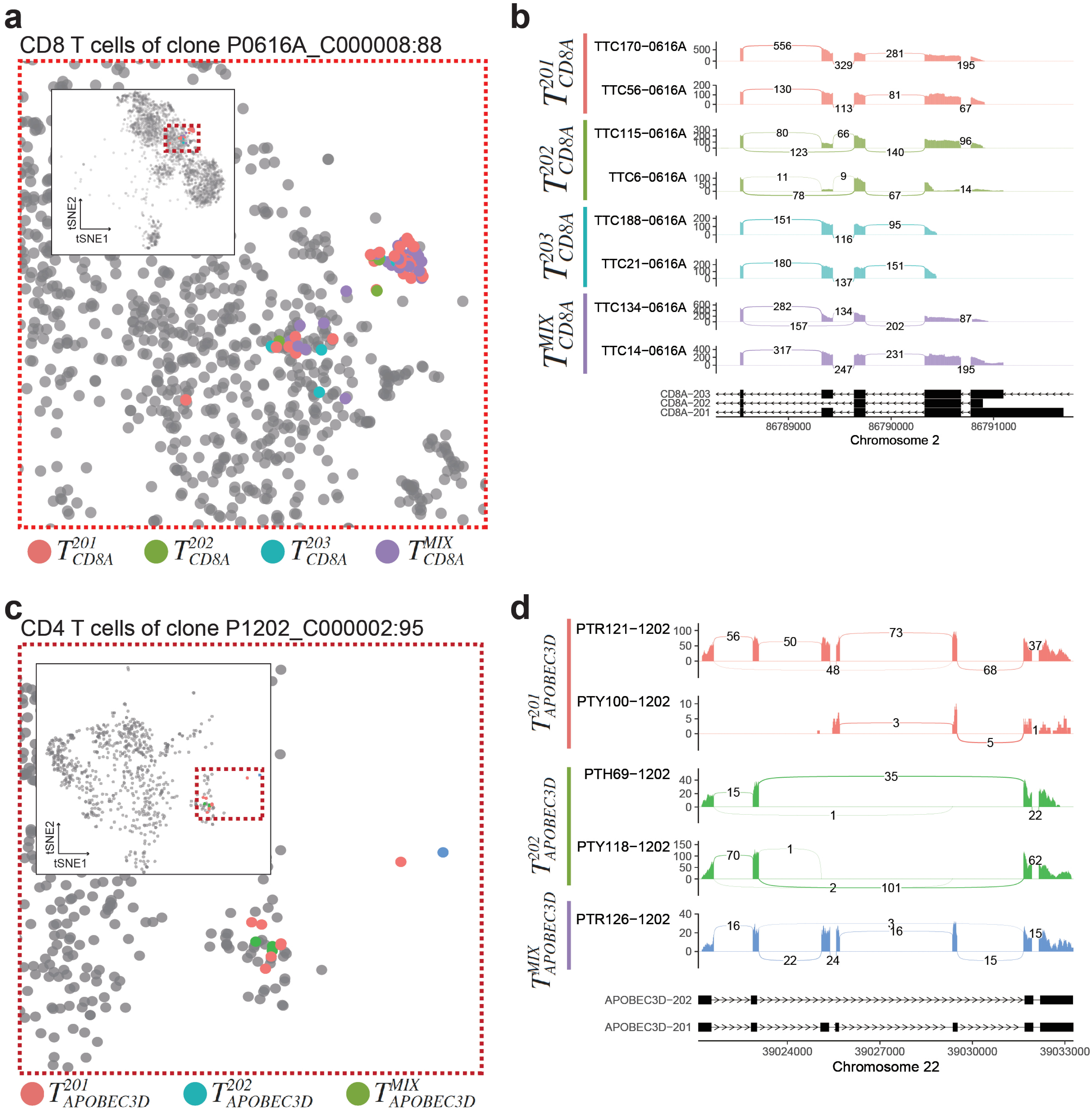
